# A unified Method for assessing the Observability of Dynamic Complex Systems

**DOI:** 10.1101/2022.01.21.477230

**Authors:** Juan G. Diaz Ochoa

## Abstract

**Problem:** Systems theory applied to biology and medicine assumes that the complexity of a system can be described by quasi-generic models to predict the behavior of many other similar systems. To this end, the aim of various research works in systems theory is to develop *inductive modeling* (based on data-intensive analysis) or *deductive modeling* (based on the deduction of mechanistic principles) to discover patterns and identify plausible correlations between past and present events, or to connect different causal relationships of interacting elements at different scales and compute mathematical predictions. Mathematical principles assume that there are constant and observable universal causal principles that apply to all biological systems. Nowadays, there are no suitable tools to assess the soundness of these universal causal principles, especially considering that organisms not only respond to environmental stimuli (and inherent processes) across multiple scales but also integrate information about and within these scales. This implies an uncontrollable degree of uncertainty.

**Methodology:** A method has been developed to detect the stability of causal processes by evaluating the information contained in the trajectories identified in a phase space. Time series patterns are analyzed using concepts from geometric information theory and persistent homology. In essence, recognizing these patterns in different time periods and evaluating their geometrically integrated information leads to the assessment of causal relationships. With this method, and together with the evaluation of persistent entropy in trajectories in relation to different individual systems, we have developed a method called *Φ-S diagram* as a complexity measure to recognize when organisms follow causal pathways leading to mechanistic responses.

**Results:** We calculated the Φ-S diagram of a deterministic dataset available in the ICU repository to test the method’s interpretability. We also calculated the Φ-S diagram of time series from health data available in the same repository. This includes patients’ physiological response to sport measured with wearables outside laboratory conditions. We confirmed the mechanistic nature of both datasets in both calculations. In addition, there is evidence that some individuals show a high degree of autonomous response and variability. Therefore, persistent individual variability may limit the ability to observe the cardiac response. In this study, we present the first demonstration of the concept of developing a more robust framework for representing complex biological systems.

## 1 Introduction

The central idea of systems biology is to reduce complex systems into interconnected elements, leading to deductive and white-box models consisting of causal paths derived from data obtained from the relevant physicochemical interactions of the system, or black-box models based on the automatic integration of large amounts of information, for example through deep learning to identify relevant system patterns (Baker et al., 2018; Zitnik et al., 2019). The hope behind this effort is to obtain all available information about the function, effects of system perturbations and future development of the biosystem. The derivation of causal pathways in such a manner considers living systems as emergent phenomena that result from interactions among different elements operating at various scales. This implies that for fixed scales it is possible to derive causal pathways and distinguish these pathways from causal mechanisms to build robust models (Baker et al., 2018; Kostić et al., 2020; Zitnik et al., 2019).

So, we face the following problem: if a model can be defined for a complex system (for example, in biology), then the complexity can be compressed in such a way that the most relevant biological characteristics are preserved, and causal pathways can be identified^2^; but if complexity is inherent, persistent, and irreducible, then it is necessary to assess the amount of complexity that could ultimately hinder the identification of principles and thus causal pathways which are required to define robust models (an idea close to the concept of complex systems biology introduced by Ma’ayan in 2017 (Ma’ayan, 2017)).

We solve this problem by calculating persistent topological features directly from data points, evaluating their geometric information, and proposing a way to measure the limits of methods for identifying causal pathways (and thus model identification). In addition, since the evaluation of topological persistence is robust against noise (Emrani et al., 2014) and the evaluation of persistent entropy allows the detection of nonlinearity across several scales (Diaz Ochoa, 2020) this represents a plausible strategy to better evaluate the possibility of identifying universal causal pathways^3^, the observability of complex systems and, in this way, the identification of robust models.

In the following section, we provide a theoretical background in systems biology to justify the development of causal pathways and geometric information. In Section 3, we review the mathematical basis for the combination of persistent topology and geometric integrated information, as well as the definition of the *Φ-S* Diagram. Thereafter, in section 4 we make an exemplary computation using deterministic datasets as well as patient data obtained from individuals using wearables measuring the heart response to acceleration (measured on three axes). Finally, in section 5 we discuss our approach and in section 6 present the main conclusions as well as an outlook for the further development of this technique.

## 2 Theoretical background: Systems theory and causality

### 2.1 Problem definition: systems biology and causality

In a systemic approach, interacting elements are analyzed in relation to system structure, context, and causal boundaries, emphasizing the emergent characteristics of the system^4^. The advantage of such systems is that complex behavior at larger scales can be characterized by understanding the interacting elements at lower scales and their context, which provides a promising way to observe and better understand the system, for example by using graph models; such systemic approaches are fundamental to the definition of systems biology and systems medicine. These systemic approaches have a clear pragmatic advantage, since they guide the identification and understanding of biological mechanisms that can be explained by mathematical models (Green, 2021)^5^ (Boogerd et al., 2007) useful in biotechnology and biomedicine for developing synthetic organisms, efficient bioprocesses, drugs, and therapies (Alon, 2006). Often, identifying these models requires a lot of effort, not only in defining the right modeling scale, equations, and parameters to describe biological processes in deductive models (Liu et al., 2013), but also in dealing with the amount of information required to build high-quality inductive models.

But such studies often ignore the fact that physical coarse-grained systems can be much more effective in terms of intrinsic cause-effect power than conventional systems based on the description of microscopic phenomena. Accordingly, *while mechanisms are generally characterized in causal terms, it is not the case that each cause acts through or is part of some mechanism, which is understood as a more or less complex arrangement of causal factors* (Reiss and Ankeny, 2016). Thus, organisms can behave as a unitary “whole” from their own intrinsic perspective to maintain homeostatic states, which implies that living systems are much more than mere “systems” with mechanistic responses, since they are autonomous (from a “coarse grained” perspective), capable of integrating self-defined and self-maintained borders, which implies *some notion of intrinsic causal control* (Marshall et al., 2017). Therefore, it is necessary to realize *the uniqueness of certain basic principles of biology that are not applicable to the inanimate world*. ^6^ This notion might include the idea that biological systems are not trivially reducible to physical laws (Walker, 2019).

The main problem with this autonomy is that a “unitary whole^7^”must account for the intervention of different elements across various scales, limiting the ability to make objective observations, as well as clear definitions of what are the appropriate “coarse-grained” rules required to correctly describe a system. Integrated system information can be characterized as a set of “coarse-grained” rules, a concept that was used to investigate the relationship between this integrated system information and the suitability of evolving systems (Joshi et al., 2013). This autonomy could imply that models in systems biology have limited applicability, a preoccupation that has been shared by many scientists and academics, i.e. the possibility that biological systems are not reducible to a Hamiltonian function (Walker, 2019). In contrast, biology is characterized by the constant absence of universal principles and branched causality, which would not be able to derive consistent and rigorous descriptions of biological systems, similar to physics or even chemistry (Ellis and Kopel, 2019).

Therefore, we are not simply dealing with network models with evolutionary links, but in general with network models whose nodes (representing internal states) and links (representing couplings between these states) change autonomously and holistically, depending on the current scale of observation, which is defined by the system itself (Diaz Ochoa, 2020). This therefore represents a type of autonomy that is not reducible (Koch, 2019), implying systems that are much more complex than dynamic and evolving networks: the representation of nodes and links is currently a “coarse-grained” state that is accessed from the outside, but determined by the system itself. Simply put, we deal with systems with irreparable inherent uncertainty. Current mathematical modelling paradigms focus on defining models to predict future outcomes. But in recent years alternative approaches have investigated the development of models that focus more on assessing persistent systems uncertainty. Thus, “models to *predict and control the future” are replaced by “models to map our ignorance about the future”* (Ravetz, 2003).

Furthermore, the problem of identification of causal laws in biological systems is still an open research issue, not only because systems show a degree of autonomy, but also because of non-linearity and multi-scaling of biological systems (according to Bizzarri et al.): *i.) Nonlinearity and multi-level feedbacks in even small systems can have unexpected consequences. A linear logic becomes inadequate because the distinction between cause and effect is lost, and the explanation for how a process occurs will require an understanding of how relationships change over time;* ii.) *The current Gene Regulatory Network (GRN) formalism is not well-suited to explain the influence of spatially distributed physical factors, most of which manifest as constraints, arising from multiple interactions and physical fields*; iii.) *Linearity is disrupted by branching pathways and unidirectionality is broken by feedback, which can occur on different levels of biological organization (at a molecular, cellular, tissue-, organ- and organism level)* (Bizzarri et al., 2019). *Thus, models of single-level interactions are broken by biological hierarchy*.

### 2.2 State of the art

#### 2.2.1 Causal analysis in timeseries: a short survey

In spite of these limitations to define well-suited causal laws in biology, a few works have sought to devise methods for determining pathways in complex systems to facilitate model identification (Baker et al., 2018), as well as to assess the degree of autonomy of a complex biological system and map our own degree of ignorance or observability of this system, i.e. our capacity to identify models and their corresponding parameters (Liu et al., 2013).

Recent research has focused on causality tests by using time series, for instance using Granger Causality (Fujita et al., 2010) or the inference of causality from bivariate time series (Paluš and Vejmelka, 2007). However, this causality inference based on the analysis of time series is currently impaired by bias in data that is possibly noisy (Paluš and Vejmelka, 2007), as well as the difficulty to capture instantaneous and non-linear causal relationships (Berzuini et al., 2012).

#### 2.2.2 Causal analysis in timeseries and topology: a novel approach

The main difference between the approach reported in this investigation and other approaches is to analyze patterns contained in time series and extract a measure of complexity from these patterns to evaluate the possibility of deriving causal paths (and not trying to derive such causal paths directly). To solve this problem, we depart from the fact that the input data contains inherent imperfections generated by the organism’s non-linearity, multiscale character, and autonomy, regardless of the quality of the sampled data. To this end we employ persistent homology, which is a tool from algebraic topology useful in situations where the “scale” is not known a priori (Edelsbrunner et al., 2000), as a kind of unsupervised machine learning identifying patterns in time series (Stolz et al., 2017), to quantify the degree of causal relationships between different time periods of different dynamic parameters using a quantity Φ_*P*_, which is essentially a measure of the integrated information of persistent topological characteristics in the time series (Oizumi et al., 2016).

Furthermore, with the same method we compute potential differences in persistent structures between different individuals from their time series. This leads to a measurement of persistent entropy *S*(Γ^*kl*^) in the population, implying low or high inter-variability of the individual’s dynamics (Diaz Ochoa, 2020). We combine both measurements, Φ_*P*_ and *S*(Γ^*kl*^), which we call *Φ-S* Diagram as a novel complexity measure, to quantify the number of persistent defects in time series that could impede the use of modeling paradigms.

## 3 Mathematical background

### 3.1 Conceptual basis

Our systems analysis is based on two main concepts: the quantification of the system’s persistent entropy (Diaz Ochoa, 2020) and the analysis of causal relationships in the system. According to the first complexity measure, persistent patterns in time series lead to individual systems’ behavior differing from the population as a whole. An analysis of this first complexity measure is intended to provide information regarding the ability to extrapolate the underlying mechanisms and theory to a large population.

The second complexity measure helps to identify the interrelationship of causal processes at different scales. It delivers information about subtle phase changes in the observed system. Analyzing the system’s autonomy extends beyond mere modeling of epistemic uncertainty^8^, as we open the possibility that this uncertainty is not fortuitous or random (or is a systematic error of observation) but is driven by the inherent autonomy of living processes. Using persistent homology, we focus our analysis on integrated information.

### 3.2 Integrated information

As complex systems are essentially non-reducible, integrated information theory has become an effective tool to measure how these processes should be understood as a whole. Fully connected systems may differ in causal processes because of different degrees of microscopic connectivity. This theory, which has its origins in neurosciences, essentially quantifies multiple causal influences among elements of a system, such that it serves as a marker about how a nervous system integrates information (Koch, 2019), which is a fundamental notion defining consciousness. However, the same theory has been used as an alternative complexity measure useful to measure causal influences among elements in different complex systems like physics, economics or biology, and has been for instance used to understand the evolution of complex organisms based on the relation between their ability to integrate information and their biological fitness (Joshi et al., 2013).

Our reasoning is to use this measure of complexity to evaluate possible changes in causal pathways. These changes could be generated by changes in intrinsic connectivity between the system’s internal states. This implies that any data analysis implicitly requires a condition of connectivity, i.e., the existence of a structure 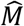 that integrates information and allows the computation of a trajectory Γ (defined in a phase space with normalized parameters), such that 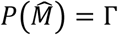. The change in these causal paths is eventually generated by environmental factors: While stable mechanisms are associated with constant causal paths (low complexity), changes in intrinsic connectivity lead to changes in causal paths (and therefore a change in measured complexity).

To this end we adapt the concept of “intrinsic cause effect power Φ” (Marshall et al., 2017) to assess the degree of autonomy of the system. According to this theory, fine-grained behavior cannot be simply reduced to interacting elements, delivering a measure of the autonomy of the system (Marshall et al., 2017) (Koch, 2019). We adapt the concept of integrated geometrical information Φ_*G*_ (Oizumi et al., 2016), considering all the possible connected and disconnected states. However, instead of analyzing the states as connected / disconnected (Marshall et al., 2017) we assume that we *“observe”* the whole state and evaluate Φ in different time periods, assuming that in such periods there are connected / disconnected internal states, which cannot be partially or fully accessed and which are encoded by the structure 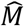.

From the geometrical integrated information theory, we consider the following three fundamental postulates:

> **<u>Postulate 1</u>**: The extent of influence is quantified by the minimal difference between models that are fully connected and those that are disconnected.
>
> **<u>Postulate 2</u>**: A disconnected model system satisfies a Markov condition.
>
> **<u>Postulate 3</u>**: The difference between a connected and a disconnected model is measured by a Kullback-Leiber (KL) divergence.

According to these postulates, and following the proof provided by Oizumi et al. (Oizumi et al., 2016)

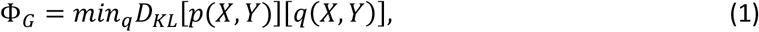

where *X* = {*x*_1_, *x*_2_, ⋯} and *Y* = {*y*_1,_ *y*_2,_ …} are the past and present states of the system, *p* is the joint probability function of connected states, and *q* is the joint probability function of disconnected states, and *D*_*KL*_ is the Kulback-Leiber entropy, defined as 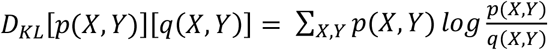

Accordingly, Φ_*G*_ is upper bounded by the mutual information *I*(*x*_*t*_′, *x*_*t*_), such that 0 ≤ Φ_*G*_ ≤ *I*(*x*_*t*_′, *x*_*t*_). Thus, Φ_*G*_ fulfills the conditions required for integrated information. Therefore, we have different scenarios:

- low information loss by mechanistic systems;
- information loss if the system behaves autonomously;
- in case of fully disconnected and aleatory systems, all information will be lost^9^.

In the case of dynamic systems, this joint probability function is characterized by the system’s trajectories in the phase space Γ, Phase space generated by connected and less connected systems is described by equation (1).

Considering that the trajectories are computed based on intrinsic information processing based on interconnected structures 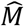, such that the computation of a trajectory can be defined as 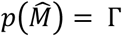, then after introducing this expression into equation 1, and based on the result of Oizumi et al. (Oizumi et al., 2016) we obtain

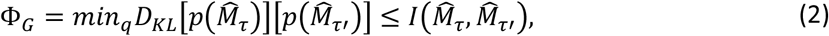

i.e., 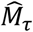 and 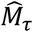 can be two different structures, where 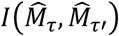 represents the mutual information. According to this definition, the divergence of Φ_*G*_ implies a divergence of microstates, which simultaneously measures the degree of autonomy of the system, because the probability distribution depending on 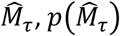, behaves in a different way as the second probability distribution, 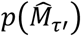, depending on the response to the environment, i.e. 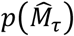 is equivalent to *p*(*x*_*t*_′, *x*_*t*_) and 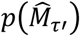 is equivalent to *q*(*x*_*t*_′, *x*_*t*_). Observe that we are comparing the system in two different periods of time, which is essentially a difference respect the original method in the eq. 1.

This approach is helpful to track possible inherent changes in 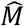. This can only be accomplished if we assume adaptation and/or evolution of the system. Thus, instead of analyzing different models leading to autonomous systems (Oizumi et al., 2016) we are analyzing the coarse grained state of the system with an underlying structure 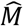. In this analysis, we assume that we do not have information about 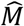; however, the analysis of the trajectory Γ provides information about the preservation of causal rules, and therefore about the autonomy of the system, and therefore about the possibility of determining 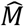 as the underlying causal structure of Γ when Φ_*G*_ → 0.

Of course, we cannot analyze connected and disconnected systems based only on Γ. But we can analyze the structure of Γ by estimating the difference of persistence bars *L*[Γ] inside different time periods, as demonstrated in previous works (Diaz Ochoa, 2020).

To this end we assume that Γ owns a topology that reflects the periodic behavior of a signal with Euler characteristics. This means Γ owns a function *g* with a compact subset of ℝ^*D*^ and 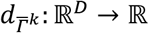 the distance function of 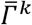. The function 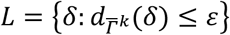 represents the set of persistent bars *δ*_*ε*_ containing the length of the topological feature extracted from the periodic behavior of the signal. In this context, a barcode is the persistence analogue of a Betti number. The *kth* Betti number of a complex, acts as a coarse numerical measure of the topological feature *H*_*k*_. Key topological features *H*_*k*_ include zero (connected points) and first order topology (loops) (Diaz Ochoa, 2020). Thus, the model estimates a hierarchical grouping of topological characteristics of higher order leading to a type of invariant represented by bar codes (Pun et al., 2018). As shown in figure 1, this methodology is used to track subtle changes in characteristic trajectories over different time periods.

**Figure 1.**
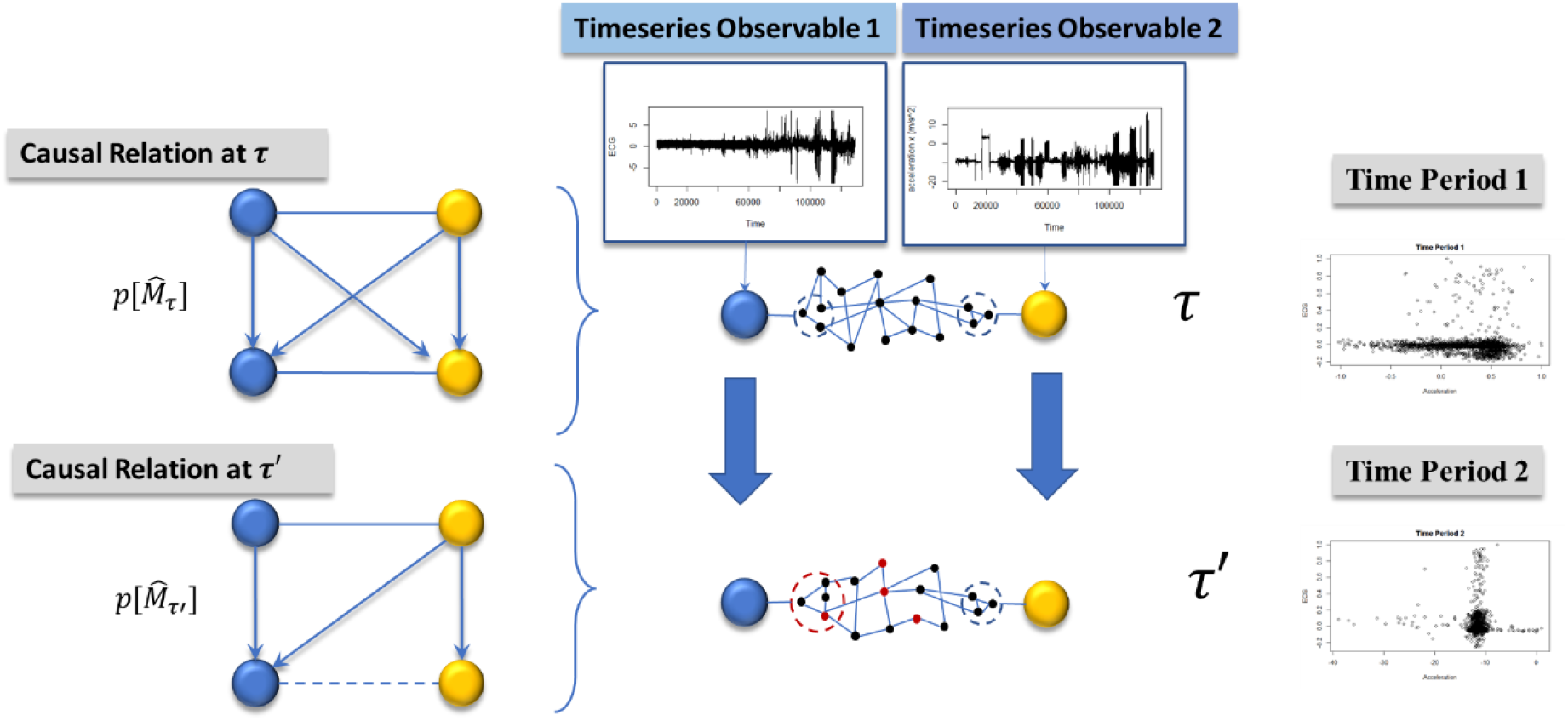
Causal relationships of a system with changing interactions (internal and external). As opposed to defining an interactome (Sanchez et al., 1999), we estimate the stability of a coarse graining approach that is necessary to define the final observations.

We compute the Kullback Leiber distance of *L*[Γ], assuming that they are implicitly generated by systems with different connectivity, such that the equation 2 gets the following form:

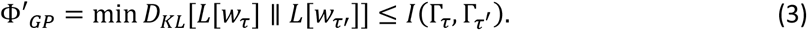

Taking into account the persistent topological characteristics of the individual organism’s trajectory, Φ_*P*_∼Φ′_*GP*_ represents the geometrically integrated information. According to these postulates, the measurement of the causal relation between underlying connectivity 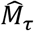 and 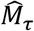 can be mathematically accounted for by the Kulback-Leiber divergence between the probability distributions of fully connected and disconnected systems ^10^, i.e., by measuring the degree of independency between the topological characteristics in different time periods. The workflow of this computation is shown in Figure 2. We used the TDA package in R for all the computations^11^.

**Figure 2.**
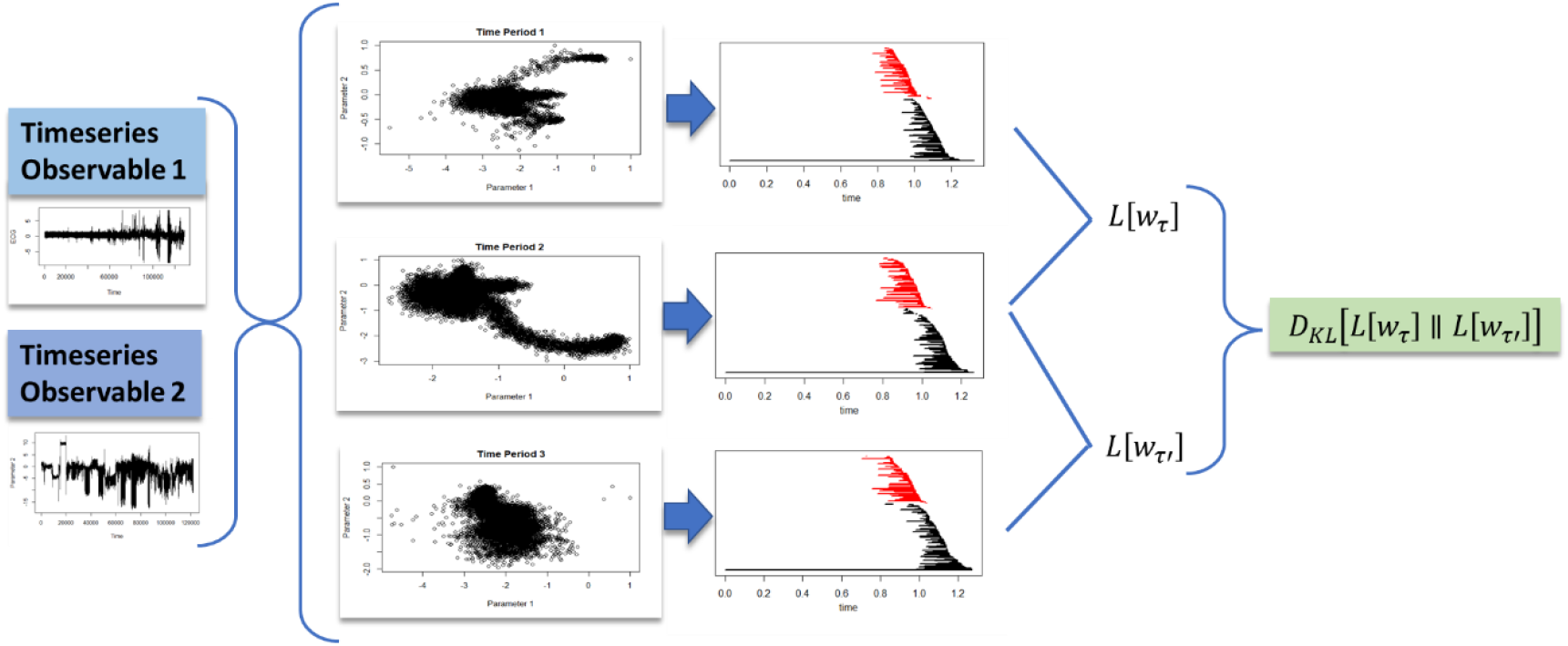
Workflow for the computation of **Φ**_**GP**_ based on the computation of persistent bars obtained from time series. This includes the construction of clouds of points from time series extracted from different time periods. In addition, it includes the calculation of persistence bars and the final calculation of Kallback-Leiber entropy for **Φ**_**GP**_ estimation.

As a result of these evaluations, it is impossible to determine whether a system has a clear causal structure derived from a mechanistic causal structure 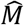. As systems explore their environment in various ways, they should be able to establish different inherent causal relations to interpret that environment; this implies that the connectome must also change accordingly. According to these postulates, if there is a deviation in the measured probability distribution, as illustrated by the workflow in figure 3, then there is probably an intrinsic change in the system (internal mechanisms as well as its interaction with the environment), impairing its observability (Figure 2), i.e., the ability to identify mechanisms interlinking different observables.

**Figure 3.**
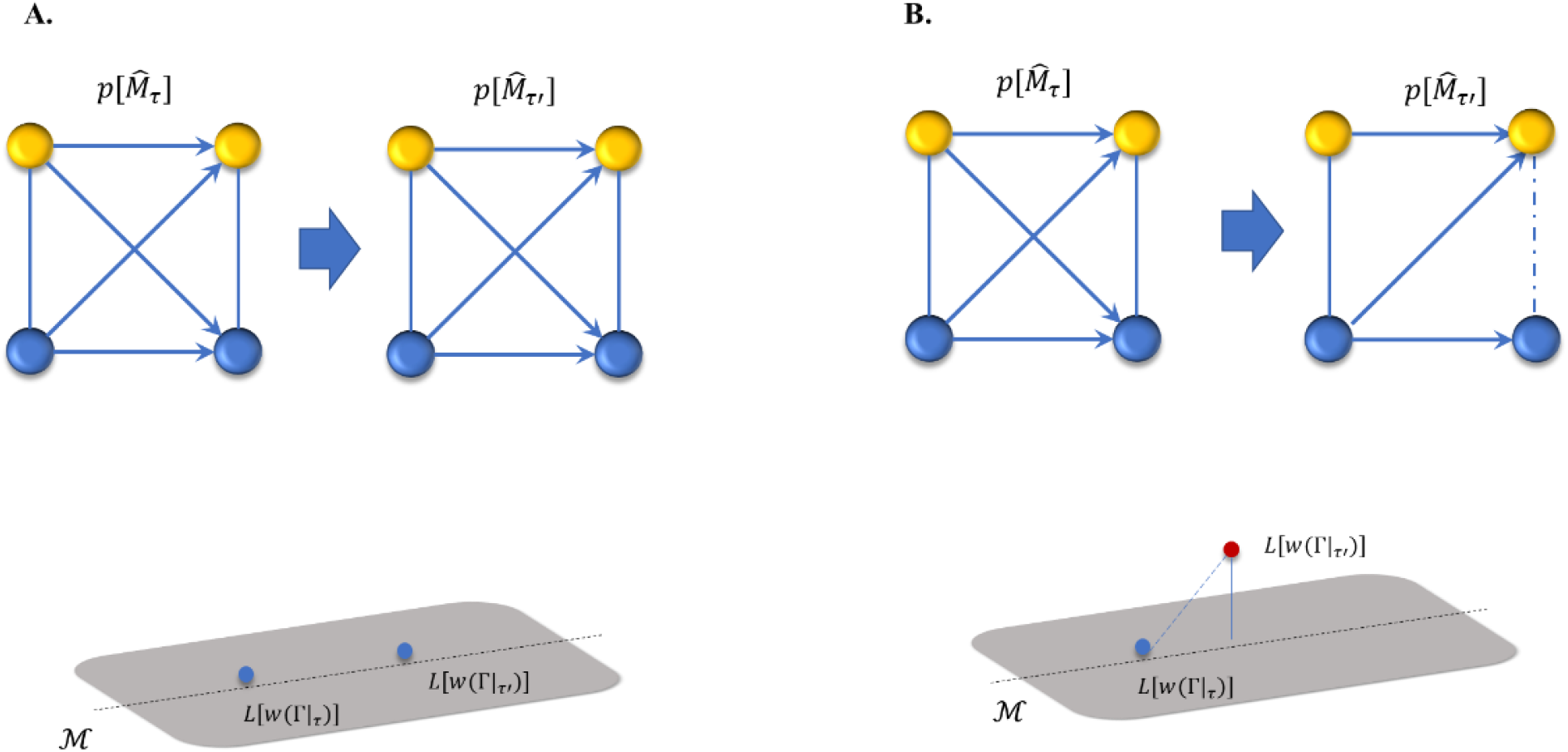
Persistence bars are computed in response to changes in internal causality in the system topology.

This situation is different from the mere concept of evolving networks since we are not only tracking a constant causal difference in a single system (dynamic and evolving system with dynamic interlinks), but also the interindividual variations, such that some individuals might evolve, while other systems possess a stable inherent structure.

The tracking of inter-species differences suggests a degree of autonomy inherent in the systems. Thus, the goal is not to model uncertainty to get a complete model^12^, but rather to learn how uncertainties can push systems to act autonomously.

According to these postulates, and the concept of mutual information^13^ (Baudot et al., 2019), the upper bound of the mutual information is the join information or join entropy (Cover and Thomas, 2006), which measures the uncertainty in the set of observations, and is defined as 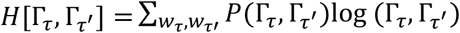, where *P*(Γ_*τ*_, Γ_*τ*′_) is the joint probability. Therefore, equation 3 can be expressed as follows:

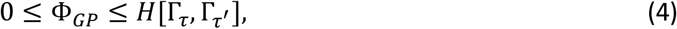

as a result, if Φ_*GP*_ → 0 then connectivity remains invariant and is constrained to a single manifold. Otherwise, the probability distribution structure diverges. Therefore, when Φ_*GP*_ → *H*[Γ_*τ*_, Γ_*τ*_′], then it implies that the inferred causal relations have the same degree of uncertainty as the observed causal relations, which eventually leads to extreme variables and aleatory systems (see Figure 3). The interpretation of this expression and its practical application is provided in the next section.

### *3.3Φ-S* Diagram

While the previous analysis aimed to assess the degree of internal connectivity between the elements of an observed system at a given scale, the estimation of persistent entropy (Diaz Ochoa, 2020) helps to assess the relative inherent differences between individual organisms. Thus, by using the measurement of geometrical (and persistent) integrated information as well as persistent entropy we can deploy a diagram to estimate the regions where different organisms probably behave in an individual and autonomous way.

The *Φ-S Diagram* provides an answer to the following questions: i) can the underlying mechanisms and theory be extrapolated to another organism or population? and ii) are these mechanisms obeying constant causal relationships?

The positive answer to both questions is the basis for universal theories, as well as models that can be extrapolated to any other system or organism.

According to this representation, we define a vector Φ_*S*_ for each observed organism *k* or system ordering the amount of integrated geometric information Φ_*GP*_ (causal analysis) and the mean amount of persistent entropy *S*(Γ^*kl*^), i.e. 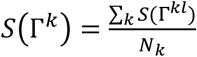, where *N*_*k*_ is the total amount of other reference organisms. Thus, the *Φ-S* Diagram for each organism *k* is defined as

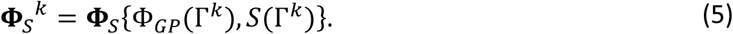

The representation of this vector and the distances measured on the main axes Φ_*GP*_ and *S*(Γ^*k*^), and the relative distance to low entropy values provides a map about observability / autonomy of the population. The interpretation of the diagram is presented in figure 3, for systems acting in a deterministic manner (Figure 4, i.) and in an autonomous manner with a high degree of individual variability (Figure 4, ii.).

**Figure 4.**
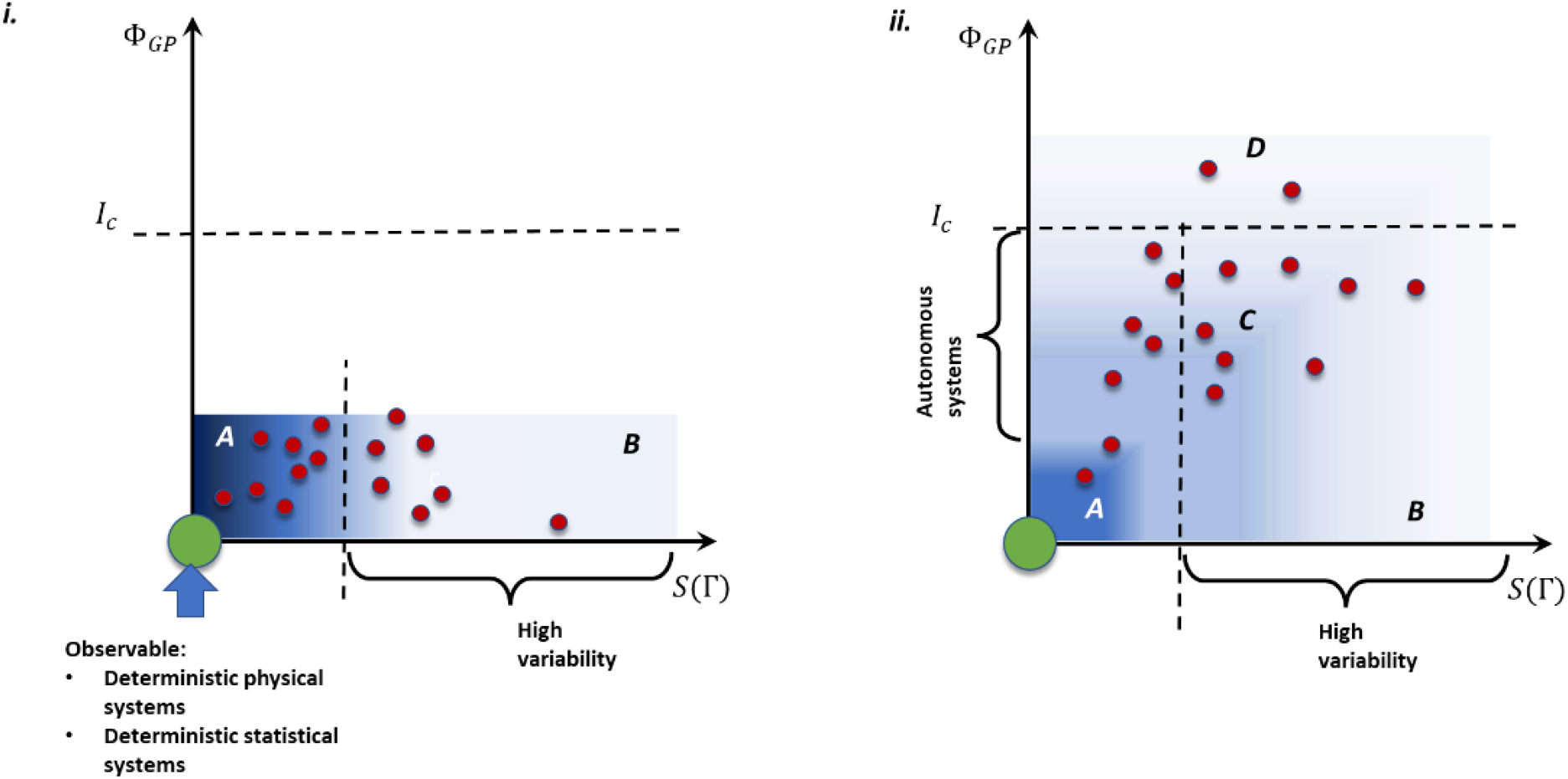
Interpretation of the Φ-S Diagram based on the composition of the geometric integrated information **Φ**_**GP**_ and the persistent entropy, for mechanistic (region A), mechanistic but high intersystem variability (region B), autonomous (region C) and aleatory systems (region D).

The short persistent bars in the diagram essentially represent noise and are therefore filtered in the final evaluation, such that any entropy measurement indicates the relative grade of disorder of persistent patterns in the paths, and not the inherent noise of the system. The vector **Φ**_*S*_^*k*^ represents the degree of disorder in persistent patterns derived from the measured paths, while *S*(Γ^*k*^) represents the relative degree of disorder of these paths (degree of individual behavior) and Φ_*GP*_ (Γ^*k*^) represents the degree of autonomy.

The usability of the *Φ-S Diagram* depends on an approximate definition of threshold values defining the three regions represented in Figure 4. Here we assume that the degree of autonomy corresponds to states changing between different regular individual mechanisms characterized by Φ_*S*_ → 0, and completely chaotic states characterized by Φ_*S*_ ≫ 0 (aleatory system). In order to better understand this diagram, we set the following criteria to determine whether the system is mechanistic, autonomous, or aleatory:

1. If we consider the probability that half of the persistent patterns are mutually divergent, then we can replace this probability in the definition of entropy to obtain *S*(Γ^*kl*^) ≈ 0.67. This approximation helps to define a threshold to find whether the system’s states mutually diverge or not: below this value the persistent patterns tend to be non-divergent (similar mechanisms across different systems, region A in figure 4); otherwise, patterns are divergent, representing high intersystem variability (region B, figure 4).
2. If Φ_*GP*_ lies below the mean value of the joint entropy, and the mean joint entropy is relatively low, then the system behaves mechanistically; otherwise, there are several phase transitions in the system, and it is likely to be autonomous based on its information processing (region C, figure 4), or aleatory (region D, figure 4).

Of course, we don’t value the system’s autonomy in the same way as the integrated information, since in principle we do not have access to the system’s structure. However, Φ_*S*_ indirectly reveals the coarsegrained autonomy of the system: if 0 < Φ_*GP*_ < *H*[Γ_*τ*_, Γ_*τ*_′], then this indicates that there are phase changes in the trajectories that do not necessarily originate from autonomous behavior. In addition, interindividual variability indicates that these phase changes are individual and, therefore, autonomous in nature.

With these two threshold values, it is then possible to fix the space where simple causal – mechanistic principles can be established. In general, when ∥**Φ**_*S*_∥ → 0 then stable inherent causal structures are responsible for mechanisms leading to model identification. Otherwise, it is possible to establish a probable non-observability (rooted in coarse-grained causality).

This diagram is like navigating on a map, helping to assess the degree of observability of single organisms within a population. In both cases, if both entropies are zero, the system is probably observable and can be described using mathematical models. Once this is guaranteed, assess whether the system’s inherent mechanisms can be represented by simply black box (or white box) models. Using this approach, it is possible to determine whether an empirical approach is the most appropriate method of understanding the system.

## 4 Results

We need benchmark data to understand how the diagram in figure 4 works, with both a deterministic component (to test the computation of Φ_*GP*_ (Γ^*k*^)) and an aleatory component.

We first tested this methodology on a deterministic dataset consisting of combining semi-periodic time series which are *highly periodic, but never repeat themselves*^14^. This data is fully synthetic and has no physical units. In order to create different individual datasets, different columns of the original data were selected and permuted per individual dataset (see supplementary information for a detailed explanation of this dataset). Thus, we intend to obtain a deterministic model that leads to the identification of models and their parameters. In addition, we also want a high degree of intraindividual variability since the original time series are permuted to generate each individual system. This should be characterized by a value **Φ**_*S*_^*k*^that represents both a high degree of causality and a relatively low persistence of defects (accordingly, it is expected that this data can be characterized by region B of the *Φ-S Diagram* in figure 4).

The analysis for three different periods is deployed in Figure 5, where the distribution of [**Φ**_*S*_^*k*^]_*i*_ for each individual *i*, is represented as a distribution. From the computation of **Φ**_*S*_^*k*^ we have found, as expected, that almost for all the systems Φ_*G*_ < 0.5, i.e., there is a high probability to share same intrinsic causal rules (low autonomy). Simultaneously we observe a high inter-individual variability, i.e., *S*(Γ^*kl*^) > 0. This implies that there are persistent characteristics that differ between individual systems. Thus, this benchmark data corroborates the basic functionality of the *Φ-S Diagram*.

**Figure 5.**
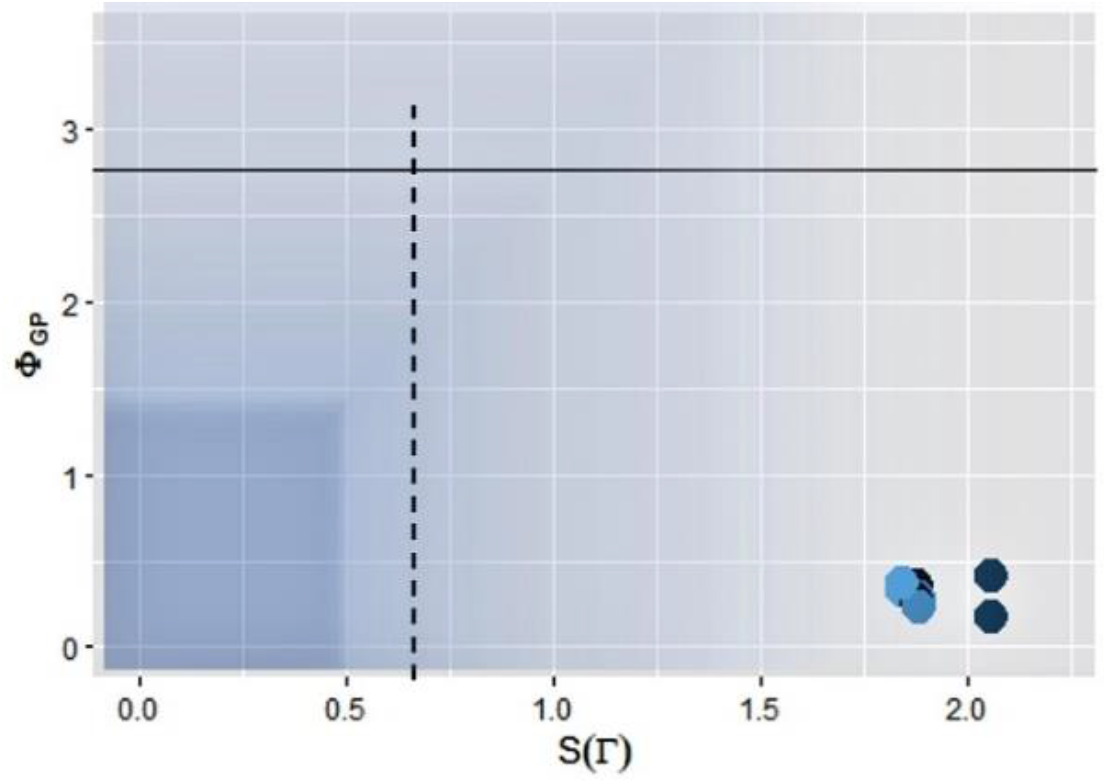
Φ-S Diagram of the benchark data

In our next example we use the M-Health data^15^ (Banos et al., 2014) to analyze the correlation between cardiac response to exercise (see appendix 1). In this database individuals were provided with wearables to measure acceleration in three axes and other physiological parameters like ECG to measure the response of the participants to sport. This data was obtained in the framework of a project to develop methods integrating mHealth data. To this end, the participants in the experiment wear a sensor positioned on their chest providing 2-lead ECG measurements and providing basic heart monitoring. Exercise sensors are accelerometers measuring the individual’s acceleration (in *m*/*s*^2^) placed on the subject’s chest, right wrist and left ankle and attached using elastic straps (see the supplementary information for more detailed information about this dataset). The purpose of this study is to measure the response to sport in real-life environments outside the laboratory.

*The heart rate regulation system was conceptualized as a complex network, with non-linear feedforward and feedback inputs. These data exhibit chaotic and non-linear dynamics as a result of interactions between physiological oscillators, functional state changes, and noise* (Dimitriev et al., 2020; Voss et al., 2009).

Despite this non-linear character, we assume that it should be possible to generate a simple *A* ⟷ *B* model based on the correlation between both states, i.e. *A*(*x, y, z*) = *f*_1_(*B*), and *B* = *f*_2_(*A*(*x, y, z*)) where *A*(*x, y, z*) is the acceleration measured in three different axes, and *B* is the heart response measured using an electrocardiogram (figure 6 A). This simple model implies that for different individuals in a small population there is a high probability to find similar causal relationships in the response to exercise, i.e., there is a straightforward and mechanistic causal relation between the heart response and physical exercise. Since we are analyzing along the tree axis, we can first analyze Φ_*G*_ as a 3D distribution^16^.

**Figure 6.**
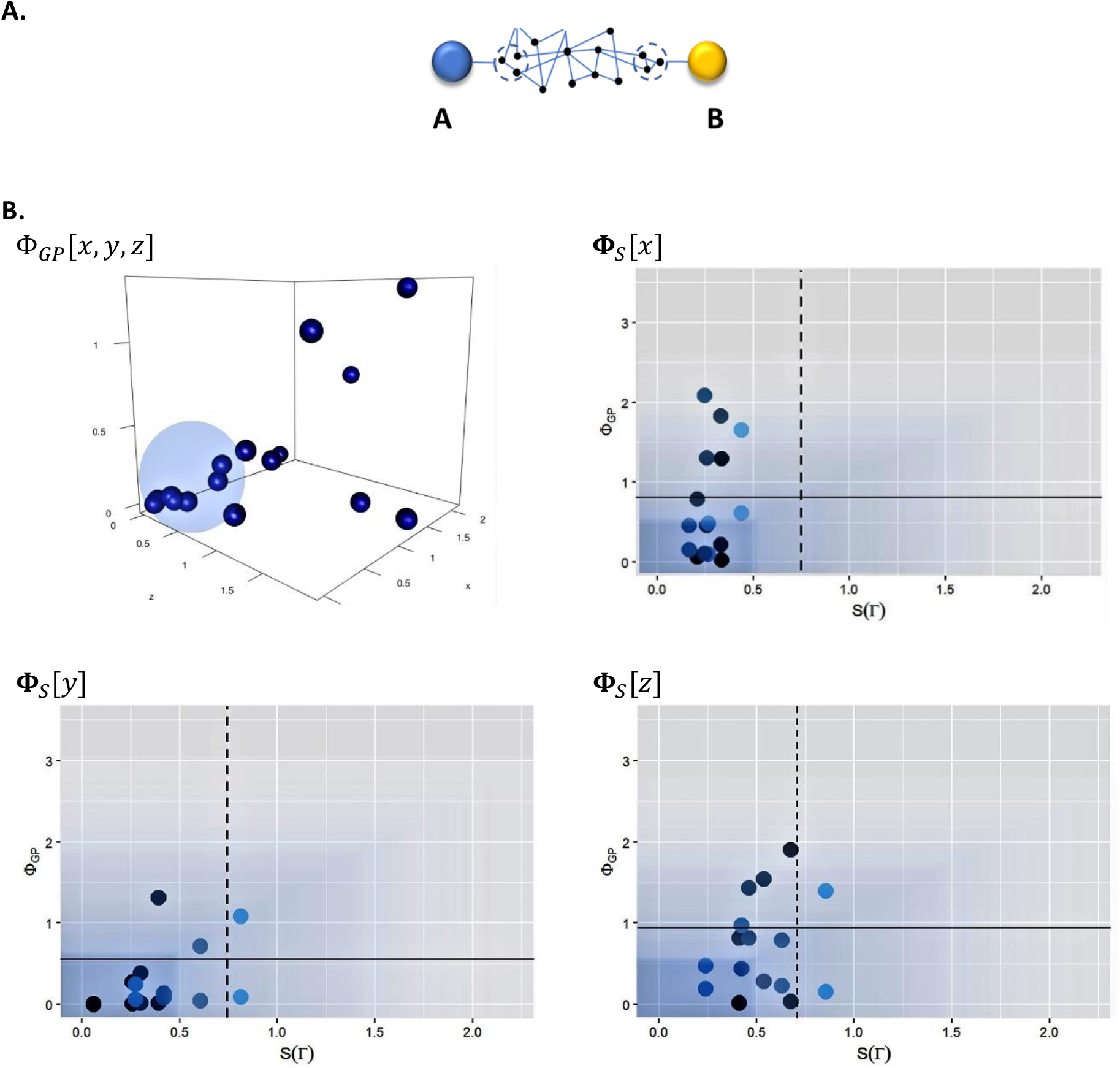
A: Based on the assumption that physical activity (measured as acceleration A) correlates with cardiac response (measured on B). Figure B: computaiton of Φ_GP [x,y,z] and Φ_S Diagram for each acceleration axis. The possibility to induce/deduce a model requires a low causal entropy; however, a first inspection of this entropy along the three axes demonstrates that only a few individuals have this low causal entropy (Figure B, 3D diagram).

To understand the meaning of the present analysis, the 3D representation can be helpful: when this complexity measure “condenses” inside the blue bubble, the causal relations are stable, indicating that this system can be modelled, that is, the recognized patterns can be used as features for inductive (artificial intelligence) models. In addition, there is “informational evaporation”, close to the condensation region, where it is likely that the system is autonomous. For larger “informational evaporation” the system is either highly autonomous or aleatory.

As a result, we found a de facto coupling between cardiac response and physical exercise. This is close to the blue region and confirms the feasibility of identifying both inductive and deductive models, or a combination of both. Although there are strong deviations (regions C and D in diagram 4) that suggest that the coupling is much more complex than merely chaotic.

**Φ**_*S*_^*k*^ values on the x axis indicate different movement patterns in some individuals, which suggests an autonomous response to exercise. Also, there is a high level of interindividual variability, i.e., only a few individuals can be correctly modeled using a type of universal model. Because only a few individuals fall within the blue square reserved for almost all mechanistic systems, we conclude that any modelling effort will be faced with a high degree of uncertainty, and that a common calibrated model for all individuals is feasible but will not provide adequate accuracy for all individuals (particularly for a model only on the y axis).

All the reported computations are reported in the following repository: https://github.com/2001Odisea/Phi_S_Diag_Persistent

## 5 Discussion

Many scholars have recently observed that we are currently amid a Copernican revolution in biology: like celestial mechanics, where the trajectory of the planets has been correctly described using basic physical laws represented in an elegant mathematical way, biological systems could soon also be represented with simple laws and consistent mathematical concepts. Reductionists believe biological laws should also be derived from basic physical laws (Kesić, 2016). The game of life is the most obvious example of how the combination of deterministic laws and chance, resembling random mutations and natural selection, could lead to cellular automata models mimicking simple living forms (Berto and Tagliabue, 2021)^17^. *The cojoining of two mean characteristics, namely dependence and autonomy, is mediated between extreme forms of dualism, which reject micro-dependence of some entities, and reductionism, which rejects macro-autonomy*^18^ (O’Connor, 2020), *a characteristic observed in several complex systems like complex networks*.

Such notion of emergence is the cornerstone for the formulation of systemic approaches leading to mathematical descriptions at different scales of molecules and pathways as well as whole populations of organisms, leading to an understanding of how living organisms metabolize, replicate and form populations and ecologies^19^. A systemic theory can then be derived from these simple laws, aiming to model any bio-medical system with relatively high precision, allowing one to perform synthetic biology and manipulate biological processes. In order to eradicate diseases and optimize production processes in biotechnology, modify biological processes, and even bring humans to other planets, these ideas have become a ray of hope (Verseux et al., 2016).

However, these ideas assume that causal relationships and relative uniformity between different organisms within a tolerance range can be identified. This leads to the identification of mechanisms or correlations. On the other hand, organisms sometimes exhibit high biological variability, perhaps originating not simply in the stochastics of the system, but also in their inherent macro autonomy and ability to integrate information linked to organism evolution and complexity (Joshi et al., 2013). For example, and contrary to the generally accepted notion of the prevalence of blind and random mutations that act as the engine of evolution, it has recently been discovered that mutations in *Arabidopsis Thaliana* are biased, that genes protect relevant parts of the genome, and that mutations occur less frequently in *functionally restricted regions* (Monroe et al., 2022).

Mechanistic approaches also inspired concepts that denied complex forms of autonomy in apparent simple organisms. This led to perceptions such as viewing insects as unconscious and robotic, with barely more emotional depth than lumps of stone^20^. Only recently has there been a different focus on complex biological systems^21^ who have even recognized the possibility that honeybees have behaviors that could reflect mood states (Bateson et al., 2011).

All these examples suggest that living systems are not blind at any scale. Furthermore, complex feedback loops at different scales make the identification of causal pathways very difficult (Bizzarri et al., 2019). Thus, a reductionist and systemic approach (leading to a synthetic organism) could be a limited perspective, either for an entire population or for single individuals, making necessary to identify complex forms of autonomy before models are formulated and validated.

This investigation developed a method and a complexity measure that allows to derive causal pathways from time series. This would enable us to observe the complex system, i.e., develop a model and identify its corresponding parameters. Using techniques inspired by integrated information, as a way to measure the complexity of a system in relation to its ability to integrate information(Koch, 2020)(Oizumi et al., 2016) we have created a novel methodology based on the estimation of persistent topology to analyze patterns in time series of states that would potentially be coupled through a model. Scanning patterns contained in a system’s response enables accurate assessment of the feasibility of a model and, in general, of a systemic reduction. This scanning is implemented in what we call *Φ-S Diagram*, which is essentially the combination of two measures of complexity: the integrated information observed from empirical measurements and persistent entropy.

This method assumes that systems behave as a whole, with non-trivial causal pathways (behaving as autonomous observers evaluating their environment regardless of the system’s scale), limiting our ability to identify well-defined mechanisms. Thus, the Φ-S Diagram is like a Ptolemais map, i.e., is a cartography of organisms that can be either placed in a known world due to its mechanistic character (small corner in the diagram), vs. organisms with larger entropy generated by their inherent autonomy belonging to a larger and less explored space in the map (limited-observability).

We first derived the theoretical background and performed a test on a deterministic system and real data generated by individuals performing physical activity (data described in appendix 1). In principle, both systems should be deterministic, while patient data should reflect primordial physiological variability. Thus, the obtained results are justified since the *Φ-S Diagram* can capture the deterministic (and thus causal) character of the system, while it is also able to detect biological variability. But this is just a first proof of concept that should be tested on additional data. Moreover, this method still requires intensive computation, since the detection of patterns in data clouds based on persistent homology is computationally intensive.

The additional limitation is the need to justify this method based on the state of machine learning. Persistent entropy and the autonomy of the system imply limitations that ultimately could not be overcome with mere engineering, not only in terms of a possible limitation in the way we objectively describe the world, considering that other organisms also interpret the world from a subjective perspective, but also how possible ethical problems might arise, by trying to derive objective observations. This could have serious implications for machine learning and artificial intelligence used to model biological systems. For example, *some argue that machine learning could be used to overcome the current scalability limitations of mechanistic modeling, while mechanistic models of machine learning algorithms could be used both as transient inputs and as validation frameworks* (Baker et al., 2018). Our approach assesses the feasibility of this marriage in light of the fact that systems eventually transcend beyond their expected mechanics. Specifically, *machine learning applications support statistical or correlation studies that bypass causality and focus exclusively on prediction* (Baker et al., 2018).

Thus, with the *Φ-S Diagram* we want to gain attention for a more critical evaluation of the causal background required for the derivation of any model, and the lack of founded knowledge in this causality (Berzuini et al., 2012), a problem that has been framed in the concept of post-normal science and the limitation of modeling in describing complex systems (Munda, 2008): at some point there is a factual boundary where a correlation-based model is not enough to describe a system, while a persistent empirical approach is the only way to understand the actual state of a system, which is an important fact not only for biology but also for medicine in analyzing mechanistic reactions and evaluating to what extent these causal relationships lead to medical outcomes (Reiss and Ankeny, 2016).

## 6 Conclusions

We have introduced a novel method based on the calculation of persistent homology and integrated information to detect patterns in dynamic trajectories. To track autonomous response patterns and non-trivial behavior in biomedical systems, this method identifies causal mechanisms and interindividual variability in dynamic trajectories. In principle, this method is a type of unsupervised machine learning method that relies on the fundamental topological characteristics of system trajectories. At the same time, this method leads to a novel complexity measurement, defined as a *Φ-S diagram*, which helps to determine the observability of a given system, i.e., the feasibility of representing this system with a deductive or inductive model.

From a theoretical point of view we have concluded that the causal relation between underlying connectivity can be measured using the Kullback-Leiber divergence between the patterns computed in different time periods, which is equivalent to the estimation of geometric integrated information (Oizumi et al., 2016), implying a measurement of a degree of autonomy of the system (Koch, 2019). Together with the computation of persistent entropy we derive the *Φ-S Diagram* as a measurement of both the implicit degree of causality and autonomy of a complex system.

Therefore, by calculating persistent entropy, as well as the integrated information contained in point clouds, we can estimate regularities in time series. The identification of deterministic behavior in a system (Figure 4) could lead to the identification of causal pathways. To demonstrate the applicability of this method, we selected two datasets that, in principle, can be described mechanistically: one based on pure deterministic data and the second based on physiological data captured from patients performing physical exercise. The mechanistic character of the first dataset is confirmed in diagram *Φ-S* in Figure 5; but for physiological data, and after calculating the corresponding *Φ-S* diagram in Figure 6, we obtain partial confirmation of the mechanistic cardiac response to exercise, showing that this method can detect individual physiologic responses that could deviate from a generalist model. Therefore, this first proof of concept suggests that the *Φ-S* diagram works as expected. As a result, we believe that this method can also be used to make an observability-cartography of any complex system, not just in biomedicine.

As presented, the results are preliminary and further testing and analysis is needed. In addition, this methodology will be extended to include high-dimensional datasets and other dynamic systems. Furthermore, additional work is needed to align the presented theory with the foundations and theoretical background of theoretical biology and systems biology, as well as to integrate it into a productive framework, for example in assessing how reliable it is to make decisions based on models, and in applying cartographic methods to the analysis of the degree of uncertainty of complex systems. In spite of this, we would like to point out that a new kind of cartography is required for understanding systems that have implications for decision-making, prior to defining and calibrating models.

## Competing interest

The author declares no competing interests.

## Acknowledgements

This work wouldn’t be possible without the deep discussions with Elena Ramirez about the limitations of mathematical modelling in economics, and in particular in behavioral economics. The author would like to thank two anonymous referees for their feedback, which helped clarify the ideas presented in this article.

## Supplementary information

### deterministic dataset

This dataset, called Pseudo Periodic Synthetic Time Series Data Set, can be download from this repository:

https://archive.ics.uci.edu/ml/datasets/Pseudo+Periodic+Synthetic+Time+Series

This data appears *highly periodic, but never exactly repeats itself*. The data is stored in one ASCII file. There are 10 columns, 100,000 rows. All data points are in the range -0.5 to +0.5. Rows are separated by carriage returns, columns by spaces.

Since the implemented method search for constant patterns of generated by two observables, observable 1 and 2 in a single system (figure 2), we generated synthetic data by randomly combing the different columns to generate a synthetic system; for instance, system 1 consisted of columns 1 and 5; system 2 consisted of columns 3 and 9, etc. In this way we generated 8 different systems. For the computation of the trajectory Γ in the phase space we combined two observables, Γ(*O*_1,_ *O*_2_), where *O*_*k*_ is the *k*′*th* observable, and considered normalized observables *N*_*O*_*k*_(*t*), i.e. 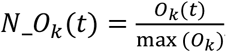, where *O*_*k*_(*t*) is the observable *k* and max (*O*_*k*_) is the maximal value of the sample of all the values of the observable *k*.

### M-Health dataset

According to the information provided by the repository, *the MHEALTH (Mobile HEALTH) dataset comprises body motion and vital signs recordings for ten volunteers of diverse profile while performing several physical activities. Sensors placed on the subject’s chest, right wrist and left ankle are used to measure the motion experienced by diverse body parts, namely, acceleration, rate of turn and magnetic field orientation. The sensor positioned on the chest also provides 2-lead ECG measurements, which can be potentially used for basic heart monitoring, checking for various arrhythmias or looking at the effects of exercise on the ECG*. This data can be downloaded from:

https://archive.ics.uci.edu/ml/datasets/MHEALTH+Dataset

An example of an ECG, as a function of time, of one of the patients is presented in the figure below.

**Figure A-1.**
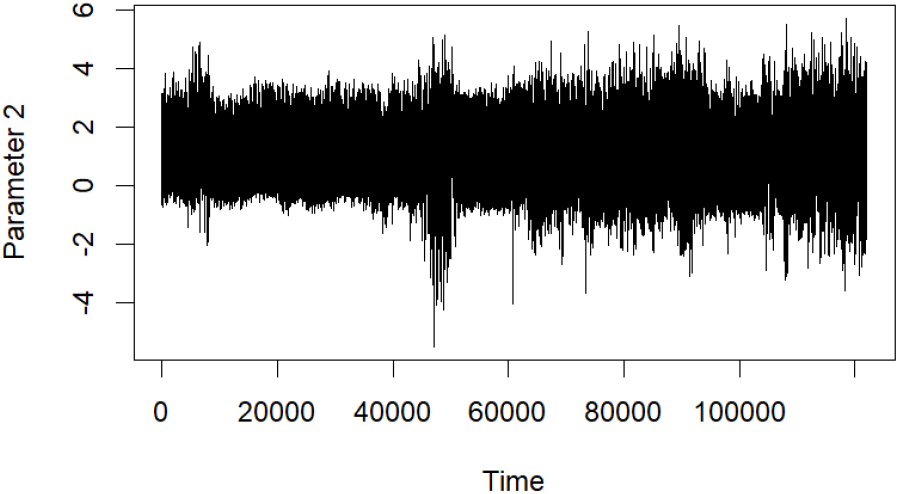
Measured ECG (Parameter 2) of patient 1 as function of time. According to the authors of this study, all sensing modalities are recorded at a sampling rate of 50 Hz.

The activity set of the participants is listed below:

L1: Standing still (1 min)

L2: Sitting and relaxing (1 min)

L3: Lying down (1 min)

L4: Walking (1 min)

L5: Climbing stairs (1 min)

L6: Waist bends forward (20x)

L7: Frontal elevation of arms (20x)

L8: Knees bending (crouching) (20x)

L9: Cycling (1 min)

L10: Jogging (1 min)

L11: Running (1 min)

L12: Jump front & back (20x)

In the reported experiments we combined the measured ECG (signal lead 1) with the acceleration measured in the chest in *m*/*s*^2^ (columns 1 to 3 in dataset). The final trajectory used for the present analysis were defined in a phase space with normalized parameters using the same normalization method applied to the deterministic dataset: For the computation of the trajectory Γ in the phase space we combined two observables, Γ(*O*_1,_ *O*_2_), where *O*_*k*_ is the *k*′*th* observable, and considered normalized observables *N*_*O*_*k*_(*t*), i.e. 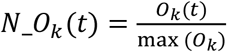, where *O*_*k*_(*t*) is the observable *k* and max (*O*_*k*_) is the maximal value of the sample of all the values of the observable *k*.

## Footnotes

2 Similar to MP3 for audio compression.

3 This led to the establishment of universal laws.

4 Using for example graph theory; for a discussion see e.g. https://plato.stanford.edu/entries/causal-models/#Grap

5 The discussion of systemic approaches can be found at https://plato.stanford.edu/entries/systems-synthetic-biology/#BiolDigiAge

6 https://www.cambridge.org/core/books/what-makes-biology-unique/autonomy-of-biology/B59719F0B2A417A7C220AD17443BC796

7 This means an entire biological unit is an organism.

8 https://towardsdatascience.com/probabilistic-neural-networks-for-breast-cancer-detection-2f1a6951e459

9 Aleatory implies complete causal disconnectedness. A stochastic uniform distribution could exhibit causal behavior if inherent mechanisms govern the trajectory of the distribution.

10 Computed in this work using the Kullback Leiber Plugin in R: https://www.rdocumentation.org/packages/entropy/versions/1.3.1/topics/KL.plugin

11 https://cran.uni-muenster.de/web/packages/TDA/vignettes/article.pdf

12 In order to achieve this goal, several modern modeling technologies have been developed.

13 A concept extracted from the topological information theory.

14 https://archive.ics.uci.edu/ml/datasets/Pseudo+Periodic+Synthetic+Time+Series

15 https://archive.ics.uci.edu/ml/datasets/MHEALTH+Dataset

16 To this end we used the sthda package in R: http://www.sthda.com/english/wiki/a-complete-guide-to-3d-visualization-device-system-in-r-r-software-and-data-visualization

17 https://plato.stanford.edu/entries/cellular-automata/#CAEmer

18 https://plato.stanford.edu/entries/properties-emergent/

19 This concept is influential in several communities, not only in biology. It biases the way living beings are considered and studied, reducing them to mere mechanisms.

20 https://www.bbc.com/future/article/20211126-why-insects-are-more-sensitive-than-they-seem

21 A feature characteristic of non-trivial autonomous systems..

## Notes

### Competing Interest Statement

The authors have declared no competing interest.

### Summary of Updates

Changes in style, language and presentation

